# Allosteric Inhibition of a Vesicular Glutamate Transporter by an Isoform-Specific Antibody

**DOI:** 10.1101/2021.06.16.447601

**Authors:** Jacob Eriksen, Fei Li, Robert M. Stroud, Robert H. Edwards

## Abstract

The role of glutamate in excitatory neurotransmission depends on its transport into synaptic vesicles by the vesicular glutamate transporters (VGLUTs). The three VGLUT isoforms exhibit a complementary distribution in the nervous system and the knockout of each produces severe, pleiotropic neurological effects. However, the available pharmacology lacks sensitivity and specificity, limiting the analysis of both transport mechanism and physiological role. To develop new molecular probes for the VGLUTs, we raised six mouse monoclonal antibodies to VGLUT2. All six bind to a structured region of VGLUT2, five to the luminal face and one to the cytosolic. Two are specific to VGLUT2 whereas the other four bind to both VGLUT1 and 2; none detect VGLUT3. Antibody 8E11 recognizes an epitope spanning the three extracellular loops in the C-domain that explains the recognition of both VGLUT1 and 2 but not VGLUT3. 8E11 also inhibits both glutamate transport and the VGLUT-associated chloride conductance. Since the antibody binds outside the substrate recognition site, it acts allosterically to inhibit function presumably by restricting conformational changes. The isoform specificity also shows that allosteric inhibition provides a mechanism to distinguish between closely related transporters.

## Introduction

The exocytic release of glutamate mediates most excitatory transmission in the mammalian nervous system. Packaging into synaptic vesicles enables glutamate release by exocytosis, and involves uptake from the cytoplasm by the vesicular glutamate transporters (VGLUTs). A proton electrochemical gradient across the synaptic vesicle membrane generated by the vacuolar-type H+-ATPase provides the driving force for vesicular glutamate transport, similar to the other vesicular neurotransmitter transporters (Anne and Gasnier, 2014; Edwards, 2007). In contrast to other vesicular neurotransmitter transporters driven by proton exchange, glutamate uptake relies on membrane potential as the main driving force (Maycox et al., 1988; Shioi, 1990), suggesting a mechanism of facilitated diffusion. However, additional ions including protons and chloride regulate vesicular glutamate transport, acting allosterically to coordinate flux with other events in the synaptic vesicle cycle (Eriksen et al., 2020).

Mammals express three VGLUT isoforms, VGLUT1 (SLC17A7), VGLUT2 (SLC17A6), and VGLUT3 (SLC17A8). The three transporters exhibit high sequence identity, especially in the transmembrane domains that make up the translocation pathway. They also do not seem to differ in intrinsic transport activity. However, they differ greatly in distribution, conferring distinct physiological roles (El Mestikawy et al., 2011; Fremeau et al., 2004). VGLUT1 is expressed primarily in the cortex, VGLUT2 in the diencephalon and brainstem (Fremeau et al., 2001; Herzog et al., 2001) and VGLUT3 often by neurons associated with a different transmitter (Fremeau et al., 2002; Gras et al., 2002; Seal et al., 2008).

Consistent with their different distributions, genetic inactivation of each VGLUT isoform produces profound but distinct neurological effects (Fremeau et al., 2004; Moechars et al., 2006; Seal et al., 2008; Wallen-Mackenzie et al., 2006; Wojcik et al., 2004). However, acute inactivation has not been possible due to the lack of specific inhibitors. The relatively low apparent affinity for glutamate (Km 1-3 mM) has presumably contributed to the difficulty identifying high affinity VGLUT inhibitors. Indeed, glutamate analogues such as aminocyclopentane-1,2-dicarboxylate and methylated forms of glutamate inhibit the VGLUTs with high micromolar potency, only slightly higher affinity than glutamate itself (Moriyama and Yamamoto, 1995; Winter and Ueda, 1993). Through unclear mechanisms, several azo dyes including Evans Blue, Chicago Sky Blue and Brilliant Yellow inhibit much more potently (Roseth et al., 1995; Tamura et al., 2014), in the low nanomolar range but lack specificity for the VGLUTs (Cao et al., 2014) and cell permeability (Pietrancosta et al., 2020). All of these compounds act as competitive inhibitors, and high homology within the binding pockets presumably accounts for the lack of specificity for VGLUT isoform. More recently, two nanobodies were raised against VGLUT1 and these recognize the cytoplasmic face of the transporter (Schenck et al., 2017). They also inhibit glutamate transport but the isoform specificity and mechanism of inhibition are not known. Here we describe the production of several antibodies to VGLUT2 that distinguish among the isoforms and show that one inhibits transport through an allosteric mechanism.

## Materials and Methods

### Antibodies

Monoclonal antibodies (mAb) against N- and C-terminally truncated rat VGLUT2 were produced as previously described (Li et al., 2020) with VGLUT2 proteoliposomes as antigen injected into mice at the Vaccine and Gene Therapy Institute (VGTI) Monoclonal Antibody Core of Oregon Health Sciences University. The antibody variable domains of hybridomas 8E11 and 9C6 were sequenced by Genscript.

### Immunofluorescence

#### Fixed cells

HEK293T cells at 90-95% confluency in 6-well plates were transfected with 2 μg pIRES2-EGFP rat pmVGLUT1, 2 or 3 (Uniprot entries: Q62634, Q9JI12 and Q7TSF2) with 6 μl FugeneHD according to the manufacturer’s protocol. The next day, the cells were seeded onto 12 mm glass coverslips coated with poly-L-lysine (1 mg/ml) in 24 well plates. One day later, the cells were washed in PBS and fixed in PBS containing 4% PFA for 30 minutes on ice. After fixation, the cells were washed 3 times in PBS before blocking and permeabilization in PBS with 5% goat serum and 0.1% Triton X-100 for 30 minutes. The cells were incubated with hybridoma supernatants diluted 1:20 in blocking buffer for 1 hour at room temperature, washed 3 times for 10 min each in PBS, incubated with goat anti-mouse antibody conjugated to Alexa Fluor 647 (1:500) in blocking buffer for 30 minutes at room temperature, washed three times in PBS for 10 minutes each, and the coverslips mounted using VECTASHIELD® Antifade Mounting Medium.

#### Live antibody feeding

Live HEK293T cells were incubated with hybridoma supernatants diluted 1:20 in full growth media for 30 minutes at RT, washed three times for 5 min each in PBS before fixation, blocking and permeabilization as above. The samples were then incubated with goat anti-mouse antibody conjugated to Alexa Fluor 647, washed and mounted as above.

#### Imaging

The specimens were imaged using a Nikon Ti inverted microscope equipped with CSU-22 spinning disk confocal and a Plan Apo VC 60x/1.4 oil objective. Images were processed using ImageJ.

### Glutamate efflux

HEK293T cells were transfected with pIRES2-EGFP pmVGLUT2 or pIRES2-EGFP as a control using Fugene HD as described above. The next day, the cells were moved into 24-well plates coated with poly-L-lysine (0.1mg/ml). Another day later, the cells were rinsed twice in Ringer’s solution (145 mM NaCl, 10 mM glucose, 10 mM HEPES – pH 7.4, 4 mM KCl, 2 mM CaCl_2_, 1 mM MgCl_2_) before loading with 25 μM ^3^H-glutamate (2 μCi) in 200 μl Ringer’s solution without or with 8E11 (1:100) for 15 minutes at 37° C. The cells were then washed three times in ice-cold Ringer’s solution containing 0.5 mM aspartate. Glutamate efflux was measured in 200 μl warm Ringer’s solution or Ringer’s solution at pH 5.5 (145 mM NaCl, 10 mM glucose, 10 mM MES – pH 5.5, 4 mM KCl, 2 mM CaCl_2_, 1 mM MgCl_2_) with 0.5 mM aspartate for 5 minutes at 37° C. 150 μl of the medium was collected and the radioactivity measured by scintillation counting. The efflux was analysed by normalizing the efflux cpm to pH 7.4 + 8E11.

### Oocyte Recordings

Recordings were performed as described previously (Eriksen et al., 2016). Briefly, *Xenopus laevis* oocytes were obtained from Ecocyte, injected with 50 ng pmVGLUT2-HA cRNA and incubated in ND96 (96 mM NaCl, 2 mM KCl, 1.8 mM CaCl_2_, 1 mM MgCl_2_, 5 mM HEPES [pH 7.4]) with 50 μg/ml tetracycline and gentamicin at 16° C for 5 days. The oocyte currents were recorded by standard two-electrode voltage clamp in Ca^2+-^free ND96 and ND96 pH 5.5 (96 mM NaCl, 2 mM KCl, 1 mM MgCl_2_, 5 mM MES [pH 5.5]). Steady-state current/voltage (I-V) relations were obtained using a protocol of 300-ms voltage steps from −120 to +60 in 10 mV steps from a holding potential of −30 mV. After first running the step protocol at pH 7.4 and pH 5.5 without antibody, the 2 μl 8E11 or control EAT-4 antibody 8A9 were added to the recording chamber, the oocytes incubated for 10 minutes, excess antibody removed by perfusion in ND96 pH 7.4 before recording at pH 7.4 and pH 5.5.

## Results

To generate probes capable of recognizing a structured domain of the VGLUTs, we injected mice with purified rat VGLUT2 reconstituted into liposomes (Li et al., 2020). The immunized mice yielded multiple monoclonal antibody hybridomas. Using supernatant from the hybridomas, we tested their binding to HEK293T cells expressing an internalization-defective, plasma membrane-targeted version of VGLUT2 (pmVGLUT2) (Eriksen et al., 2016). All six of the tested antibodies show binding to fixed, permeabilized cells expressing pmVGLUT2 (identified by the coexpressed EGFP), whereas non-transfected (EGFP-) cells in the confluent cultures show no binding (Figure 1A).

**Figure 1.**
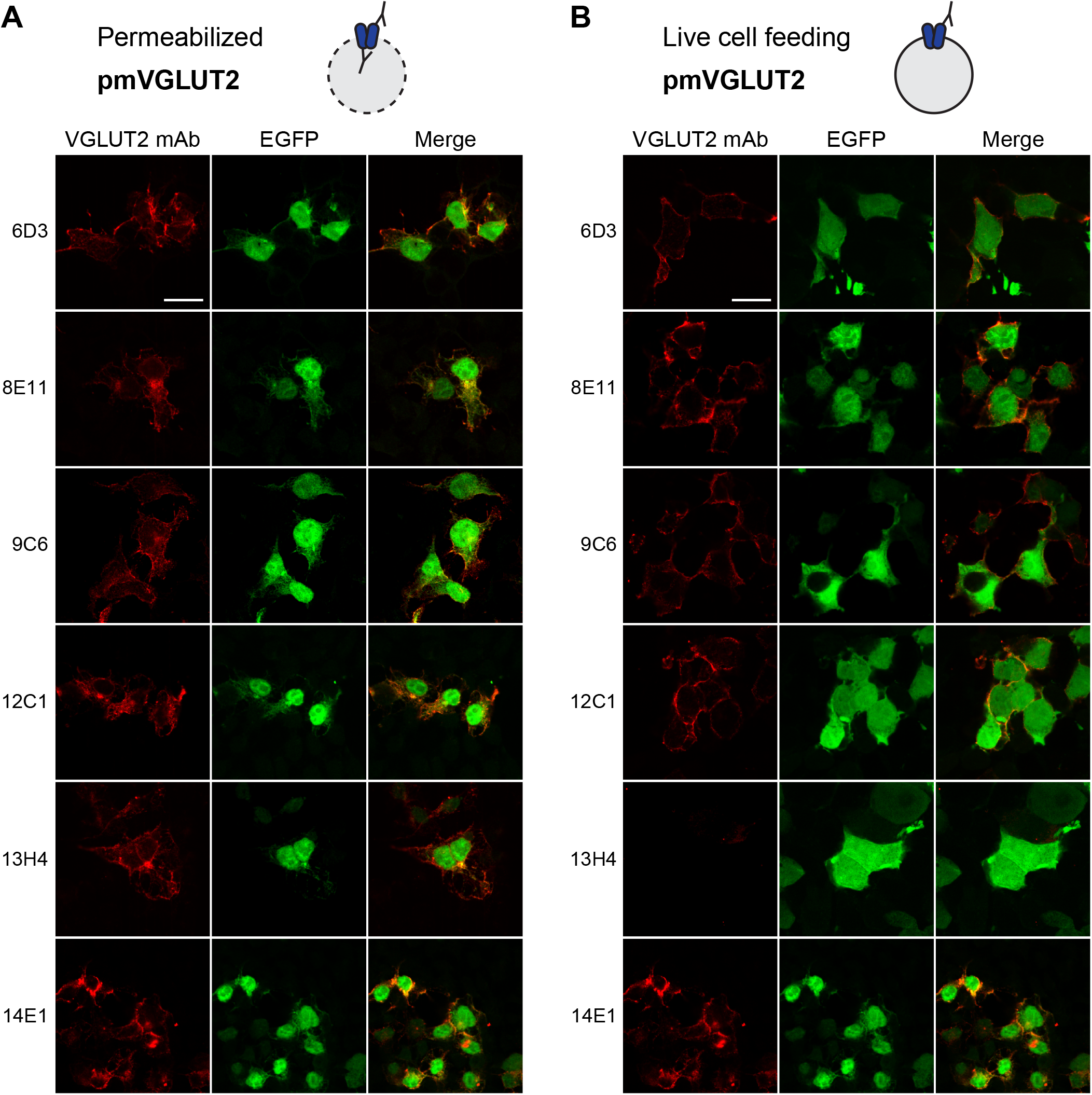
Novel monoclonal antibodies bind VGLUT2 from different sides of the membrane. HEK293T cells co-expressing pmVGLUT2 and EGFP were stained with VGLUT2 mAb hybridoma supernatants. (A) After fixation and permeabilization, all VGLUT2 mAbs label only transfected cells identified with EGFP. (B) Live HEK293T cells were incubated with hybridoma supernatants. 6D3, 8E11, 9C6, 12C1, and 14E1 but not 13H4 bind to intact cells expressing pmVGLUT2. 13H4 thus binds to the cytoplasmic face of VGLUT2. Scale bar, 20 μm.

The use of pmVGLUT2 enabled us to determine whether the antibodies bind to either the luminal or cytoplasmic face of the transporter. Live cells expressing pmVGLUT2 can only be stained by antibodies that recognize the luminal/external face of VGLUT2 whereas antibodies recognizing the cytoplasmic face will not have access to this site in cells with an intact plasma membrane (Figure 1B). Supernatants from hybridomas 6D3, 8E11, 9C6, 12C1 and 14E1 all give rise to a fluorescent signal in HEK293T cells expressing pmVGLUT2 (Figure 1B), showing that these antibodies bind to the luminal surface of VGLUT2. In contrast, 13H4, which shows specific binding to VGLUT2 in fixed, permeabilized cells, does not show any signal with intact cells. Thus, 13H4 binds to the cytoplasmic face of VGLUT2, similar to the nanobodies recently reported (Schenck et al., 2017).

We then assessed the isoform specificity of the antibodies by immunostaining cells expressing plasma membrane-targeted versions of VGLUT1 or VGLUT3. Supernatants from hybridomas 6D3, 8E11, 9C6 and 12C1 all bind to pmVGLUT1-expressing cells whereas 13H4 and 14E1 show no binding (Figure 2A). None of the antibodies show any binding to cells expressing pmVGLUT3 (Figure 2B). Thus, of the six antibodies recognizing VGLUT2, two are specific for VGLUT2 (13H4 and 14E1), four recognize both VGLUT1 and 2 (6D3, 8E11, 9C6 and 12C1) and none recognize VGLUT3. Five of the antibodies bind the luminal face of the transporter and only one (13H4) binds to the cytoplasmic side.

**Figure 2.**
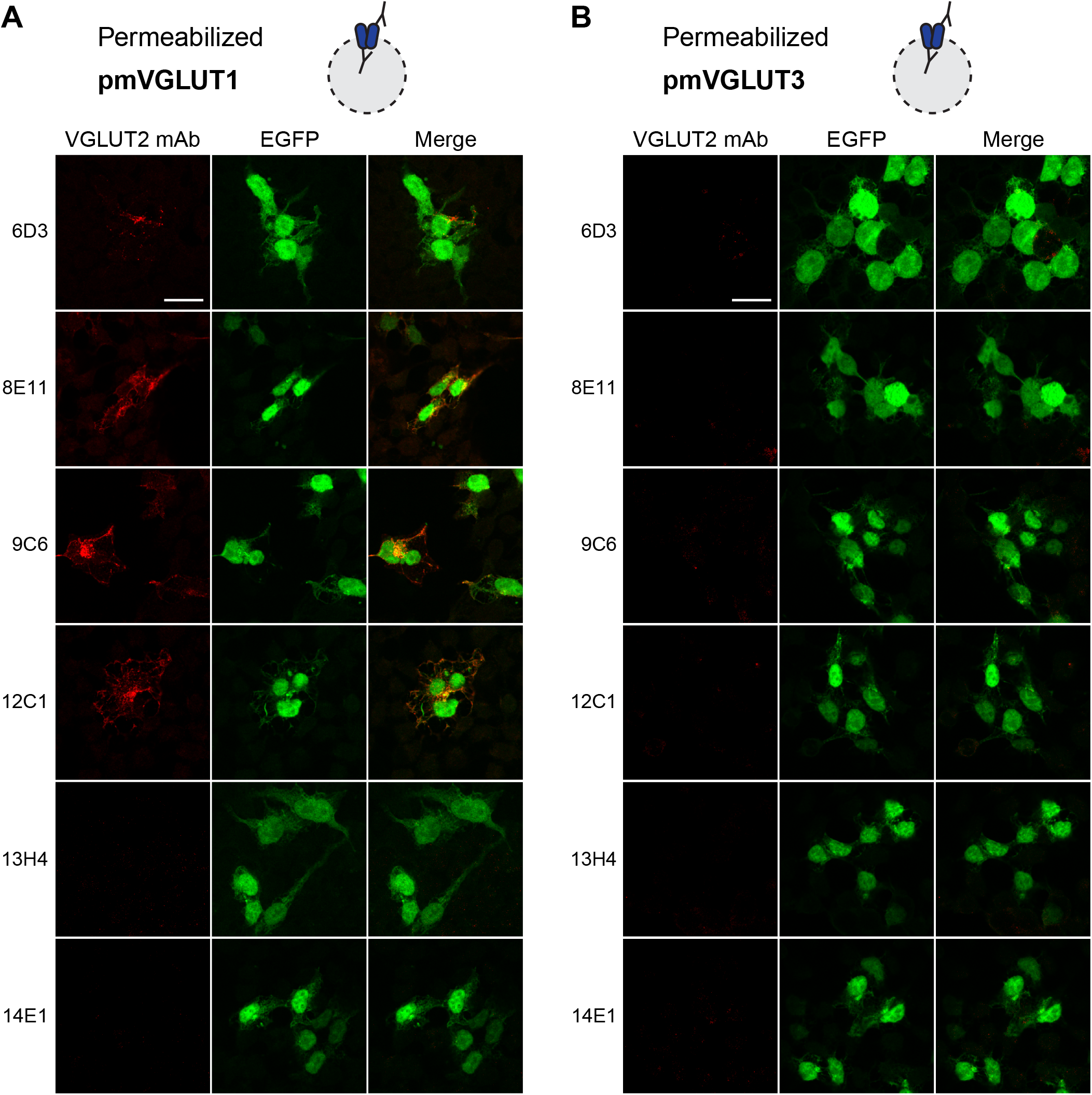
VGLUT isoform specificity of novel VGLUT2 mAbs. Fixed, permeabilized HEK293T cells expressing pmVGLUT1 (A) or pmVGLUT3 (B) were incubated with VGLUT2 mAb hybridoma supernatants as in Figure 1A. (A) mAbs 6D3, 8E11, 9C6, 12C1 bind to cells expressing pmVGLUT1, whereas 13H4 and 14E1 do not. (B) None of the mAbs bind to cells expressing pmVGLUT3. Scale bar 20 μm.

Since the antibodies were raised against folded, functional VGLUT2 (Li et al., 2020), we also determined whether they recognize a linear or structured epitope by western blotting denatured extract from cells expressing a VGLUT2-EGFP fusion. In contrast to the EGFP antibody, none of the hybridoma antibodies recognize VGLUT2-EGFP by immunoblot (Figure S1). Thus, all of the hybridoma antibodies recognize a folded epitope. The properties of the mAbs are summarized in Table 1.

**Table 1.**
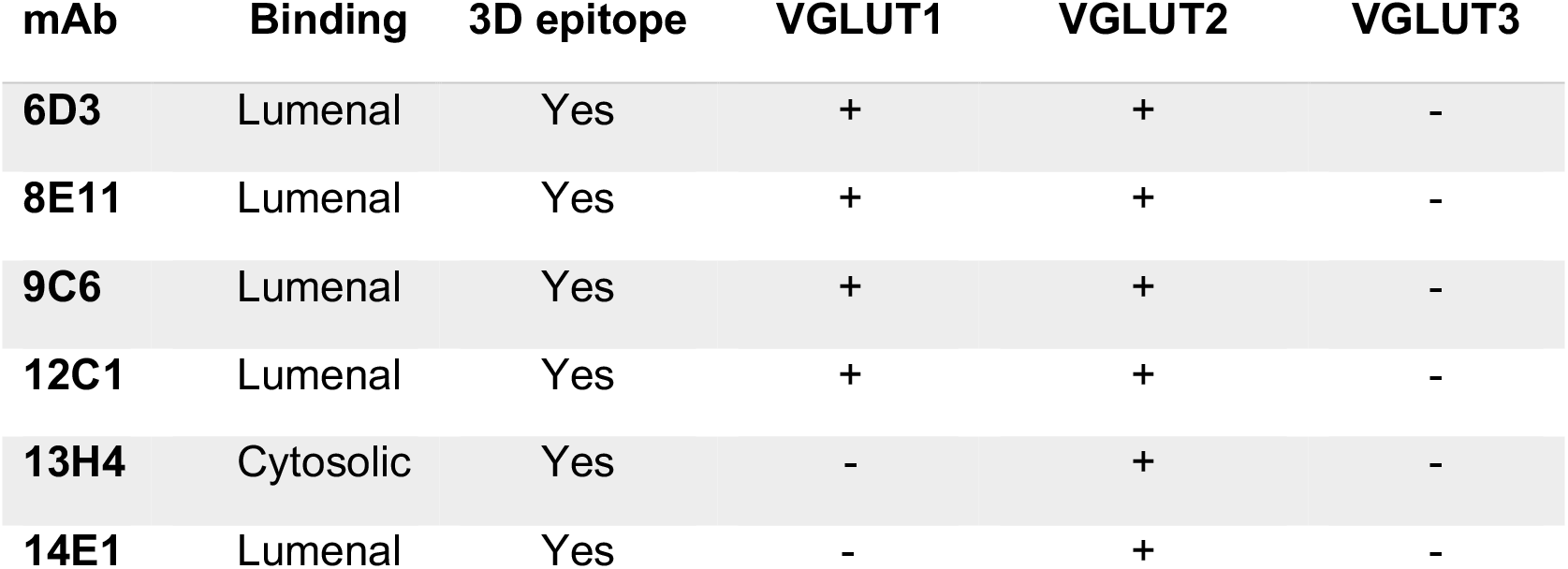
Properties of the VGLUT2 mAbs

| **mAb** | **Binding** | **3D epitope** | **VGLUT1** | **VGLUT2** | **VGLUT3** |
| --- | --- | --- | --- | --- | --- |
| **6D3** | Luminal | Yes | + | + | - |
| **8E11** | Luminal | Yes | + | + | - |
| **9C6** | Luminal | Yes | + | + | - |
| **12C1** | Luminal | Yes | + | + | - |
| **13H4** | Cytosolic | Yes | - | + | - |
| **14E1** | Luminal | Yes | - | + | - |

We previously showed that an Fab fragment of the 8E11 antibody was essential for a high resolution cryo-EM structure of VGLUT2 (Li 2020). The structure of the VGLUT2/8E11 complex showed that 8E11 binds to the lumenal side of VGLUT2, consistent with the staining of unpermeabilized cells in Figure 1B. Specifically, the Fab makes multiple polar contacts with all three short extracellular loops (ECL 4, 5, and 6) in the C-domain of the transporter (Figure 3A). In ECL 4, the backbone carbonyls of V340 and G342 in VGLUT2 hydrogen bond with the backbone amides of L102 and S52 in the 8E11 heavy chain, respectively. Also, in ECL4, the E338 sidechain forms a hydrogen bond with Y57 in the 8E11 heavy chain, and E344 with the side chains of S52 and S56 in the 8E11 heavy chain (Figure 3A). In ECL5, R409 forms a hydrogen bond with the sidechain of Y32 in the 8E11 heavy chain (Figure 3A). In ECL6, the backbone carbonyl of K471 hydrogen bonds with the sidechain of S92 in the 8E11 light chain and the sidechain of VGLUT2 R473 hydrogen bonds with Y31 and makes a salt bridge with D49 in the 8E11 light chain. Antibody 8E11 thus makes an extensive network of contacts with ECL4-6 of VGLUT2. The sequence of hybridoma antibody 9C6 is almost identical to that of 8E11 (Figure S2), suggesting a very similar interaction with VGLUT2.

**Figure 3.**
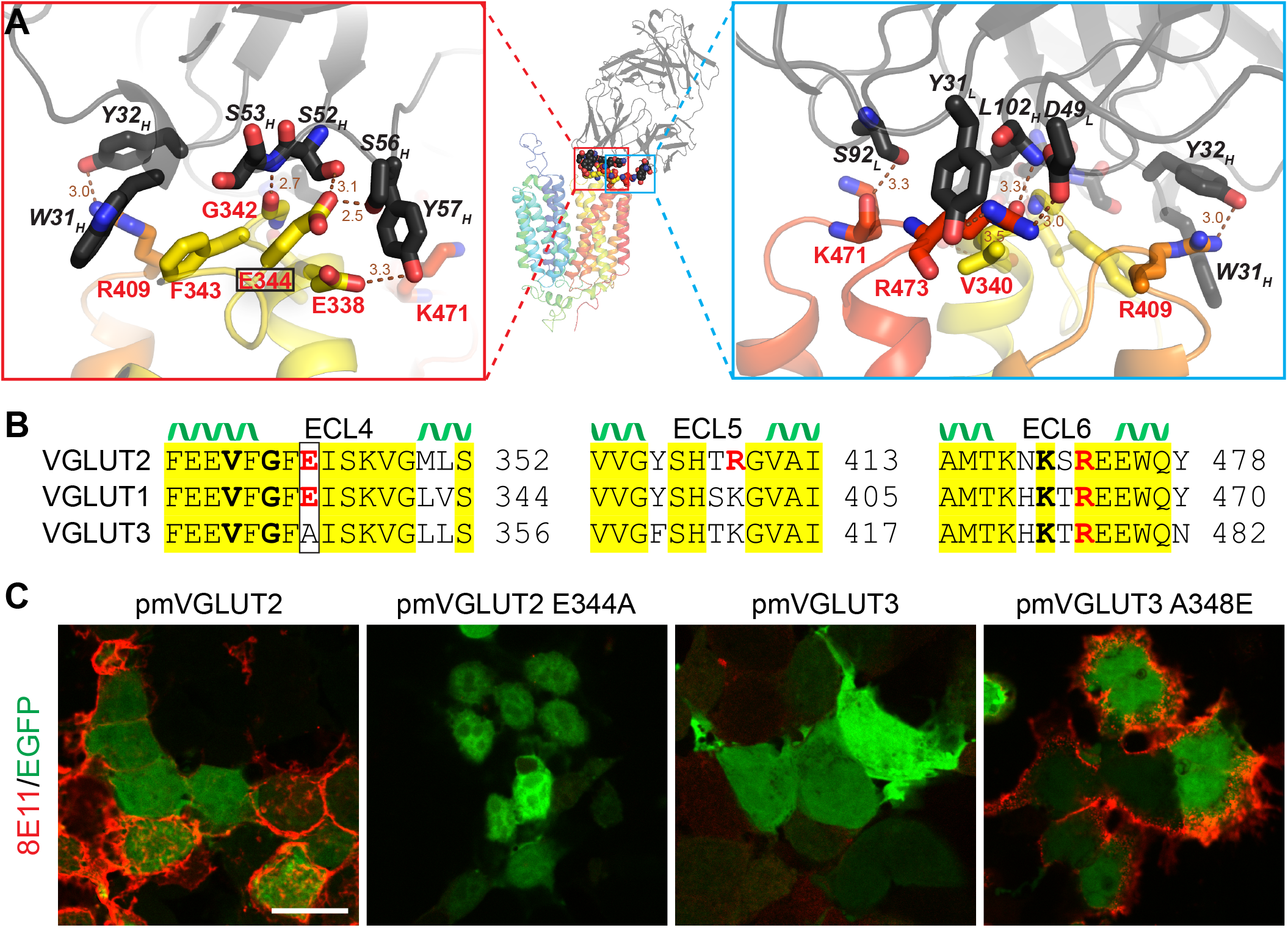
Structure of VGLUT2-8E11 reveals the basis of isoform specificity. (A) Cryo-EM structure of rat VGLUT2 (rainbow) and Fab 8E11 (black) with the interface enlarged on the left and right (PDB: 6V4D). The sequence on the right is rotated 180° around an axis perpendicular to the membrane. (B) Alignment of ECL4-6 of VGLUT1-3 with conserved residues highlighted in yellow. VGLUT residues contacting 8E11 are indicated in bold and side chain interactions in red. Glutamate 344 (boxed) is the only residue conserved in VGLUT1 and 2 but not VGLUT3. C) Live cell feeding for pmVGLUT2, pmVGLUT2 E344A, pmVGLUT3 and pmVGLUT3 A348E confirms the requirement of 8E11 for the divergent ECL4 glutamate. Scale bar 20 μm.

To understand how 8E11 recognizes VGLUT1 and 2 but not VGLUT3, we compared the sequence of ECL4-6 from all three isoforms. E344 is the only residue in ECL4-6 conserved in both VGLUT1 and 2 but not VGLUT3, where it is replaced by an alanine (Figure 3B). To determine whether E344 is required for the high affinity binding of 8E11 to VGLUT2, we replaced this residue with alanine, resulting in a loss of recognition by 8E11 (Figure 3C). Conversely, replacing A348 in VGLUT3 with glutamate was sufficient to confer recognition by 8E11 (Figure 3C). Thus, the residue at this position in ECL4 determines isoform preference by 8E11. The sequence of VGLUT isoforms in ECL4-6 are conserved from rodent to human, predicting that 8E11 and 9C6 hybridomas can also distinguish among the human isoforms (Figure S3).

To determine whether 8E11 interferes with transport, we used an assay for glutamate efflux from HEK293T cells expressing pmVGLUT2. This assay enables the addition of 8E11 to the luminal/extracellular face of VGLUT2 simply by adding 8E11 to the assay buffer. This is not possible with a native preparation such as synaptic vesicles or even after VGLUT reconstitution into proteoliposomes, which exposes the cytoplasmic but not lumenal face of the transporters (Preobraschenski et al., 2014; Schenck et al., 2017). Cells expressing pmVGLUT2 or EGFP as control were loaded with [^3^H]-glutamate (Figure 4A): HEK293T cells accumulate glutamate independent of pmVGLUT2 expression. To mimic uptake by synaptic vesicles, we triggered VGLUT-mediated glutamate efflux by incubating the preloaded cells at low pH (Figure 4A) since lumenal/extracellular protons allosterically activate the VGLUTs (Eriksen et al., 2016). Cells expressing pmVGLUT2 show increased glutamate efflux at pH 5.5 relative to pH 7.4, and control cells show no increase in efflux at low pH (Figure 4B). The addition of 8E11 abolishes the pH-dependent pmVGLUT2-mediated efflux, with no effect of the antibody on efflux from control cells (Figure 4B).

**Figure 4.**
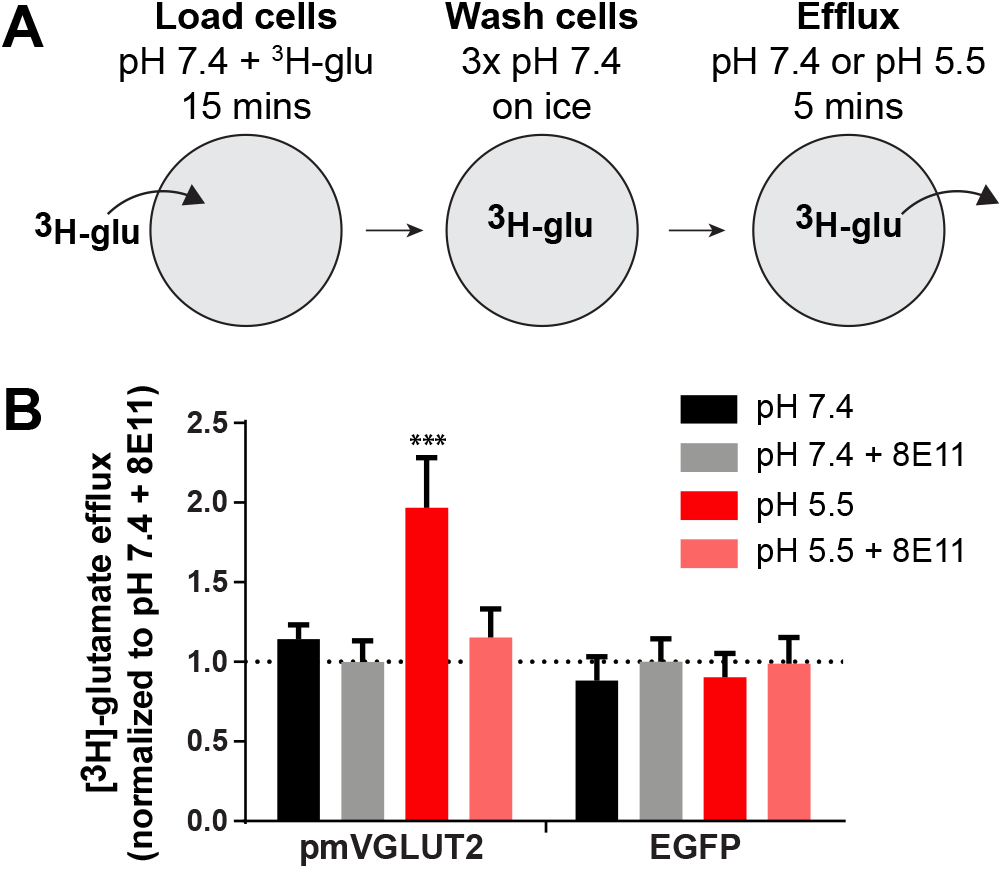
8E11 inhibits glutamate transport by VGLUT2. (A) Assay for glutamate efflux. HEK293T cells are loaded with ^3^H-glutamate in the absence or presence of 8E11, washed in cold Ringer’s solution before incubation in efflux buffer at either pH 7.4 or pH 5.5. (B) Efflux of ^3^H-glutamate from HEK293T cells expressing pmVGLUT2 + EGFP or EGFP alone. Efflux was measured with or without 8E11 at pH 7.4 and pH 5.5 (n=5). Data indicate mean ± SEM. *** p < 0.001, by two-way ANOVA.

The VGLUTs also exhibit an associated chloride conductance(Bellocchio et al., 2000; Chang et al., 2018; Eriksen et al., 2016; Schenck et al., 2009). This conductance shares many features with glutamate transport including allosteric activation by lumenal/extracellular protons and chloride, but is not coupled to glutamate flux and seems to involve a different mechanism although the two anions compete for permeation (Chang et al., 2018; Eriksen et al., 2016). To determine whether 8E11 influences the VGLUT-associated chloride conductance, we expressed pmVGLUT2 in Xenopus oocytes, where low pH triggers large VGLUT-mediated chloride currents (Figure 5A,B). Incubation of the same oocytes with 8E11 for 10 mins effectively eliminated the low pH-activated currents (90.7 ± 1.9 % inhibition of low pH-induced current). A parallel experiment with mAb directed to the C. elegans VGLUT homologue EAT-4 produced in the same way as 8E11 had no effect on the VGLUT2 chloride conductance (0.2 ± 6.5 % inhibition of low pH-induced current), demonstrating the specificity of 8E11 (Figure 5B).

**Figure 5.**
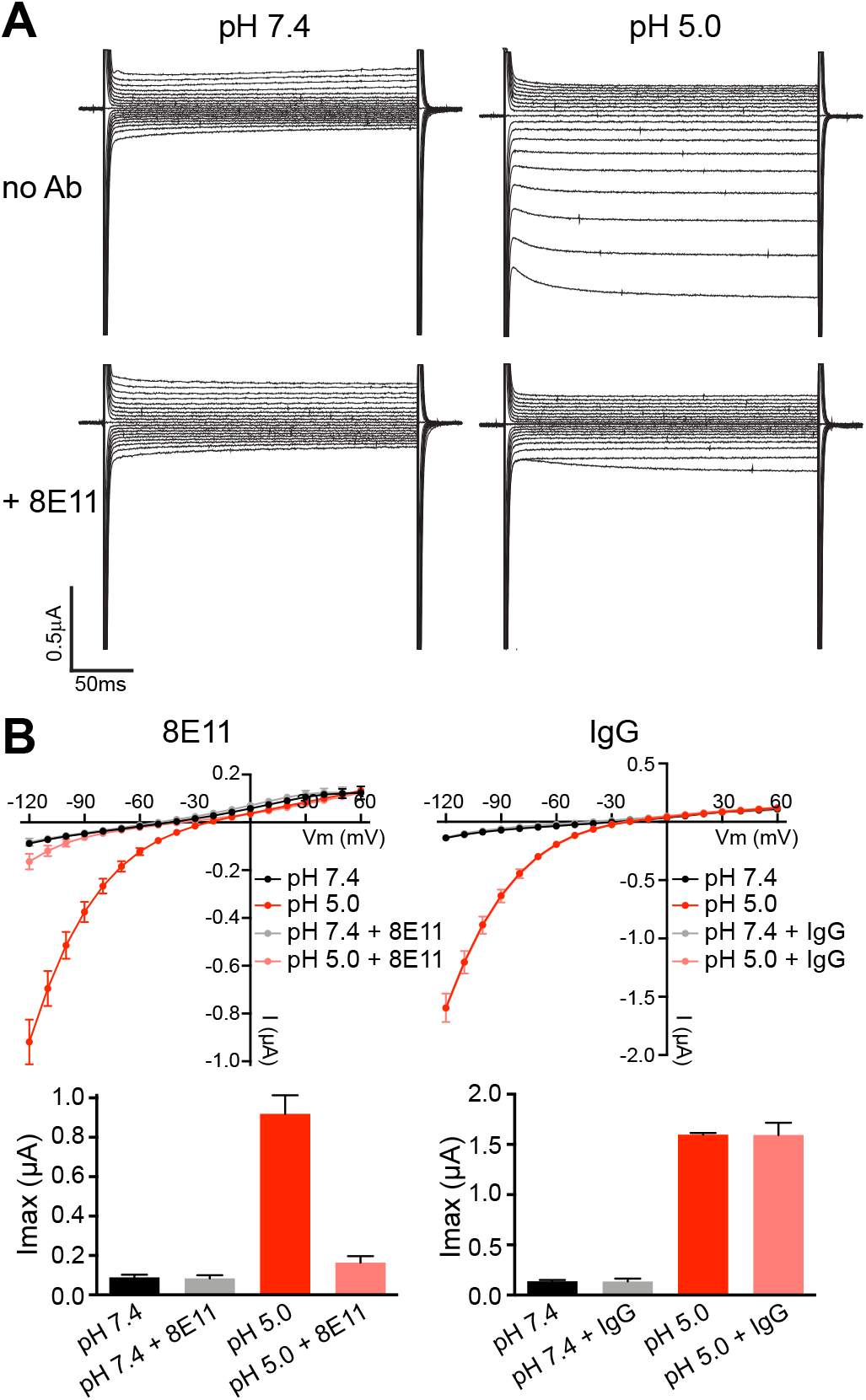
8E11 inhibits the chloride conductance associated with VGLUT2. A) Representative traces of the currents from an oocyte expressing pmVGLUT2-HA, at either pH 7.4 or 5.0, without or with 8E11. B) I-V curves of the currents from *X. laevis* oocytes expressing pmVGLUT2-HA (top) and bar graphs of the same currents at −120 mV (bottom). The currents were recorded at pH 7.4 and pH 5.5 before and after incubation with mAb 8E11 or an EAT-4 (IgG) antibody as negative control. n=7 oocytes for 8E11, n=3 oocytes for EAT-4 mAb. Data indicate mean ± SEM.

## Discussion

In contrast to the extensive and potent pharmacology for plasma membrane neurotransmitter transporters, the vesicular neurotransmitter transporters in general and the VGLUTs in particular lack potent, specific inhibitors. In this study, we characterize a series of monoclonal antibodies to the VGLUTs. Conformation-specific antibodies enable manipulation of their targets for both structural and functional studies, so we raised antibodies against a truncated form of VGLUT2 lacking part of the cytoplasmic N- and C-termini to favor the recognition of nonlinear, structured epitopes. We obtained six new antibodies that recognize structured regions of VGLUT2, including two specific for VGLUT2, four others that also bind to VGLUT1 and none that recognize VGLUT3.

Focusing on the antibody previously used to solve the structure of VGLUT2, we find that 8E11 inhibits both vesicular glutamate transport and the associated chloride conductance. In previous work, we found that glutamate transport and the associated chloride conductance undergo similar allosteric regulation by protons and chloride (Eriksen et al., 2016) [ref], and we now find that the similarity between these two activities extends to inhibition by 8E11.

The cryo-EM structure of VGLUT2 provides insight into the mechanism of inhibition by 8E11 (Li et al., 2020). The structure shows that 8E11 binds to ECL4-6 of the VGLUT2 C-domain through an extensive network of interactions. The antibody does not bind in the main cavity of the transporter where substrate is recognized, demonstrating an allosteric mode of inhibition. The requirement for 8E11 on the opposite side of the membrane from substrate makes it difficult to determine whether this allosteric form of inhibition is non-competitive. The interactions observed by cryo-EM account for the isoform specificity of 8E11, with binding to VGLUT1 and 2 but not VGLUT3. E344 in ECL4 confers binding specificity towards VGLUT1 and 2, and its replacement by alanine (as in VGLUT3) disrupts the interaction. Conversely, replacement of alanine by glutamate at this position in VGLUT3 suffices to confer binding by 8E11. Since the site of substrate recognition shows high conservation across VGLUT isoforms, binding to less well conserved sequences elsewhere in the protein confers specificity not possible with an orthosteric inhibitor. Similar to the development of allosteric modulators for receptors that distinguish among isoforms (Foster and Conn, 2017), it may thus be possible to develop inhibitors that discriminate between other closely related transporters. It would also be of considerable interest to determine how antibodies 13H4 and 14E1 distinguish between VGLUT2 and VGLUT1.

How does antibody binding at an allosteric site within the VGLUT C-domain inhibit transport activity? In general, members of the major facilitator superfamily (MFS) use a rocker-switch mechanism, where N- and C-domains achieve alternating access by a symmetrical rocking movement around the substrate binding site (Drew et al., 2021). Indeed, a comparison of the inwardly and outwardly oriented conformations of the bacterial galactonate transporter DgoT, which is closely related to the VGLUTs, shows only restricted changes within each helical bundle (Leano et al., 2019). It might therefore be expected that binding outside the main cavity to a single domain would not perturb the translocation mechanism, but 8E11 appears to trap VGLUT in a lumenal/outward-oriented conformation, which has facilitated structure determination. We propose that by binding to the three external loops, 8E11 prevents rearrangement within the C-domain required for the transition between conformations. Indeed, structural evidence from other MFS members points to asymmetric transporter intermediates even though inward open and outward open states appear symmetric (Drew et al., 2021). Like the nanobodies recently reported (Schenck et al., 2017), hybridoma 13H4 binds to the cytoplasmic face of VGLUT2 and it would be of interest to determine whether it also inhibits transport and if so, the mechanism involved.

Recently, multiple studies have used VGLUTs localized at the plasma membrane to study their diverse properties, including glutamate transport (Eriksen et al., 2016), chloride conductance (Eriksen et al., 2016; Li et al., 2020; Martineau et al., 2017) and phosphate transport (Cheret et al., 2021; Preobraschenski et al., 2018). However, the lack of potent, specific, membrane-permeant inhibitors has made it difficult to determine whether the activities monitored reflect the function of VGLUT or an endogenous protein. Our results now show that 8E11 can be used to inhibit VGLUT in these assays and to confirm that VGLUTs mediate the fluxes monitored.

## ASSOCIATED CONTENT

### Accession codes

VGLUT1 (rat): Uniprot entry Q62634, VGLUT2 (rat): Uniprot entry Q9JI12, and VGLUT3 (rat): Uniprot entry Q7TSF2

### Supporting Information

Supplementary figures S1-3

## AUTHOR INFORMATION

### Author Contributions

J.E. Performed immunostainings and VGLUT functional experiment. F.L. Prepared VGLUT for immunization, did the biochemical characterization of antibodies and determined the structure. All authors designed and analyzed experiments. J.E. and R.H.E. wrote the paper with input from the F.L. and R.M.S.

### Funding Sources

This work is supported by R01NS089713 to R.M.S. and R.H.E. R37MH50712 to R.H.E. F.L. is supported by postdoctoral fellowships from the American Heart Association (17POST33660928) and the National Institute of Mental Health (K99MH119591). The UCSF EM facility is supported by NIH grants S10OD020054 and S10OD021741. Some of this work was performed at the Stanford-SLAC Cryo-EM Center (S2C2), which is supported by the National Institutes of Health Common Fund Transformative High-Resolution Cryo-Electron Microscopy program (U24 GM129541). The content is solely the responsibility of the authors and does not necessarily represent the official views of the National Institutes of Health.

## ACKNOWLEDGEMENTS

We thank Dr. David Bulkley, Dr. Zanlin Yu, Mr. Glenn Gilbert and Mr. Matt Harrington at the UCSF cryo-EM facility and Dr. Corey Hecksel, Dr. Patrick Mitchell, Dr. Lydia-Marie Joubert and Ms. Lisa Dunn at Stanford-SLAC Cryo-EM Center (S^2^C^2^) for their support in data acquisition and computation. We thank Dr. Daniel Cawley at VGTI (OHSU) for generating the mAb and advice on working with the antibodies. We also thank Dr. Charles Craik, Dr. Markus Bohn and Dr. Koli Basu for advice and help in characterizing and working with antibodies.

## Notes

### Competing Interest Statement

The authors have declared no competing interest.

