## Supplementary Figures for "Allosteric Inhibition of a Vesicular Glutamate Transporter by an Isoform-Specific Antibody"

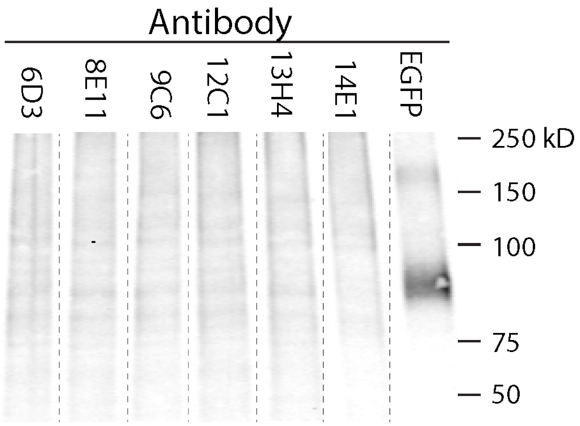


**Figure S1: VGLUT2 mAbs do not recognize denatured VGLUT2 by immunoblotting.**

Immunoblot of extracts from HEK293T cells expressing a VGLUT2-EGFP fusion. The membranes were incubated with the VGLUT2 hybridoma mAbs or EGFP antibody as control.

### Heavy chain

```
8E11  MNFGLSLIFLVLVLKGVQCEVMLVESGGGLVKPGGSLKISCAASGF 27
9C6   MNFGLSLIFLVLVLKGVQCEVMLVESGGGLVKPGGSLKISCAASGF 27
      **********************************************

8E11  TFSWYAMSWVRQTPEKRLEWVATISSAGSYTYYPDSVKGRFTISRD 73
9C6   TFSWYAMSWVRQTPEKRLEWVATISSAGSYTYYPDSVKGRFTISRD 73
      **********************************************

8E11  NAKNTLYLQMSSLRSEDTAMYYCSRVPILGRVDYWGQGTTLTVSS 118
9C6   NAKNTLYLQMSSLRSEDTAMYYCSRVPILGRVDYWGQGTTLTVSS 118
      *********************************************
```

### Light chain

```
8E11  MDFQVQIFSFLLISASVIISRGQIVLTQSPAFMSASPGEKVTMTCS 24
9C6   MDFQVQIFSFLLISASVIISRGQIVLTQSPAIMSASPGEKVTMTCS 24
      ******************************:***************

8E11  ASSSVTYMNWYQQKSGTSPKTWIYDSSRLASGVPPRFSGSGSGTSY 70
9C6   ASSSVTYMHWYQQKSGTSPKTWIYDSSRLASGVPARFSGSGSGTSY 70
      ********:*************************************

8E11  SLTISSMEAEDAATYYCQQWSSNPPIFTFGSGTKLEIK 108
9C6   SLTISSMEAEDAATYYCQHWSSNPPIFTFGSGTKLEIK 108
      ******************:*******************
```

Signal peptide-FR1-CDR1-FR2-CDR2-FR3-CDR3-FR4-

**Figure S2: mAbs 8E11 and 9C6 show sequence identity.**

Alignment of the heavy and light chain variable regions from 8E11 and 9C6. Signal peptides are shown in black, framework regions (FRs) in red and complementarity-determining regions (CDRs) in blue. Residues at the interface with VGLUT2 are shown in bold. The 8E11 and 9C6 heavy chains are 100% identical and the light chains 97% identical, with no differences at the VGLUT2/Fab interface. Numbering starts after the cleaved signal peptide.

| | ECL4 | | ECL5 | | ECL6 | |
|---|---|---|---|---|---|---|
| hVGLUT1 | FEEVFGFEISKVGLVS | 344 | VVGYSHSKGVAI | 405 | AMTKHKTREEWQY | 470 |
| mVGLUT1 | FEEVFGFEISKVGLVS | 344 | VVGYSHSKGVAI | 405 | AMTKHKTREEWQY | 470 |
| rVGLUT1 | FEEVFGFEISKVGLVS | 344 | VVGYSHSKGVAI | 405 | AMTKHKTREEWQY | 470 |
| hVGLUT2 | FEEVFGFEISKVGMLS | 352 | VVGYSHTRGVAI | 413 | AMTKNKSREEWQY | 478 |
| mVGLUT2 | FEEVFGFEISKVGMLS | 352 | VVGYSHTRGVAI | 413 | AMTKNKSREEWQY | 478 |
| rVGLUT2 | FEEVFGFEISKVGMLS | 352 | VVGYSHTRGVAI | 413 | AMTKNKSREEWQY | 478 |
| hVGLUT3 | FEEVFGFAISKVGLLS | 356 | VVGFSHTKGVAI | 417 | AMTRHKTREEWQN | 482 |
| mVGLUT3 | FEEVFGFAISKVGLLS | 369 | VVGFSHTKGVAI | 430 | AMTKHKTREEWQN | 495 |
| rVGLUT3 | FEEVFGFAISKVGLLS | 356 | VVGFSHTKGVAI | 417 | AMTKHKTREEWQN | 482 |

**Figure S3: Alignment of ECL4-6 from VGLUT1-3 of different species.**

Alignment of the 8E11 binding site in VGLUT1-3 from human, mouse and rat with conserved residues highlighted in yellow. VGLUT residues making contact with 8E11 are in bold and side chain interactions in red. Glutamate 344 (boxed) is the only non-conservative difference between VGLUT1/2 and VGLUT3.
